# *Agrobacterium*-mediated floral-dip transformation of the obligate outcrosser *Capsella grandiflora*

**DOI:** 10.1101/757328

**Authors:** Kelly Dew-Budd, Brandon David, Mark A. Beilstein

**Author notes:** Corresponding Author: Mark A. Beilstein, School of Plant Sciences – University of Arizona, P.O. Box 210036, Forbes Building, Room 303, Tucson, Arizona 85721-0036.

## Abstract

Plant transformation by floral dip has been essential for research on plant genetics. The plant family Brassicaceae is one of the most well studied plant families and contains both established and emerging genetic model species. Two emerging model species that bear on the evolution of the selfing syndrome are *Capsella grandiflora*, an obligate outcrosser, and *C. rubella*, an inbreeder. While the selfing syndrome has been well characterized at the genomic level the genetic mechanisms underlying it remain elusive, in part due to the challenges of establishing mutation lines in *C. grandiflora*. Here, we describe an efficient method for transforming *C. grandiflora* by *Agrobacterium*-mediated floral-dip while simultaneously tracking self-incompatibility loci. With the ability to transform both *C. grandiflora* and *C. rubella*, researchers have gained a valuable tool to study the progression to selfing at the genetic level.

---

The plant family Brassicaceae is a diverse group of roughly 3500 species that includes *Arabidopsis thaliana* and a number of emerging genetic model species with available whole genome sequences [1]. Functional work in many of these emerging models has been propelled by the ability to modify their genomes via *Agrobacterium*-mediated floral dip transformation. For example, transformation of Brassicaceae species has allowed for the study of leaf shape in *Cardamine hirsuta* [2], salt-tolerance in *Eutrema salsugineum* [3] and *Schrenkiella parvula* [4], and biofuel production in *Camelina sativa* [5,6]. These species are in two major linages of the Brassicaceae, suggesting a broad ability for *Agrobacterium* transformation to work in the family [7–9]. Interestingly, plants in the genus *Brassica*, a member of lineage II, are not transformable by *Agrobacterium* floral dip, but can be transformed using *Agrobacterium* in callus [10–14].

In lineage I of Brassicaceae, the congeners *Capsella grandiflora* and *C. rubella* are sister species that are 20,000 – 50,000 years diverged [15,16], but which differ in breeding system. While *C. grandiflora* is self-incompatible (SI), the SI loci responsible for this phenotype were lost or degraded in *C. rubella* resulting in self-compatibility [17]. Thus, these sister species have become important models for understanding the morphological and genomic consequences of the progression from SI to selfing. The loss of SI is associated with differences in floral morphology [18], reproductive imprinting [19], and cis-gene regulation [20,21]. To date, these differences have been examined at the genomic level but not at the functional and genetic levels because *C. grandiflora* has, to our knowledge, not previously been the subject of studies that require its transformation. This is not surprising given the added challenges presented by the introduction and maintenance of transgenes in a population of interbreeding individuals as is required for species with functional SI loci. Here, we describe a method to transform *C. grandiflora* by *Agrobacterium*-mediated floral dip, and importantly, a strategy for monitoring SI loci to efficiently maintain transgenes. These strategies will allow for the genetic dissection of phenotypic and genomic differences underlying the transition to selfing.

The *C. grandiflora* accession used in this study, 83.17, was generously provided by Stephen Wright [22]. Seeds were surface sterilized and stratified at 4°C for seven days. After germination on MS plates, seedlings were transplanted into soil and grown under a 16 hour-photoperiod at 20°C and ∼30% relative humidity.

The T-DNA cassette used for *Agrobacterium* transformation was designed to express EGFP-tagged oleosin, a seed oil body protein, under the native oleosin promoter, for easy visual screening of T1 seeds [23], and Basta resistance to facilitate screening of propagated plants post germination (Figure 1A). The T-DNA cassette also included a CRISPR construct containing SpCas9-D10A under the AtRPS5A promoter, a constitutive Arabidopsis ribosomal protein promoter, and two single-guide RNAs. The plasmid was transformed into *Agrobacterium tumefaciens* GV3101 via electroporation and uptake was confirmed by PCR amplification of the Basta resistance gene. *Agrobacterium* carrying the transgene was used to inoculate 3 ml of lysogeny broth (LB) containing the antibiotics spectinomycin and gentamycin, and incubated at 28°C for 24 hours. Following incubation, the culture was used to inoculate 500 ml of LB containing the same antibiotics. Inoculated LB was grown at 28°C for 16 hours or until it reached an optical density between 1.0 – 1.4. The *Agrobacterium* culture was spun at 4500 g, and the resulting pellet was resuspended in 500 ml of infiltration medium. Infiltration medium contains 25 g sucrose, 500 µl Gamborg’s vitamins (Bioworld), 5 µl 6-BA (10 µg/µl, Fisher), 100 µL silwet, and ddH_2_O to a final volume of 500 ml.

**Fig. 1.**
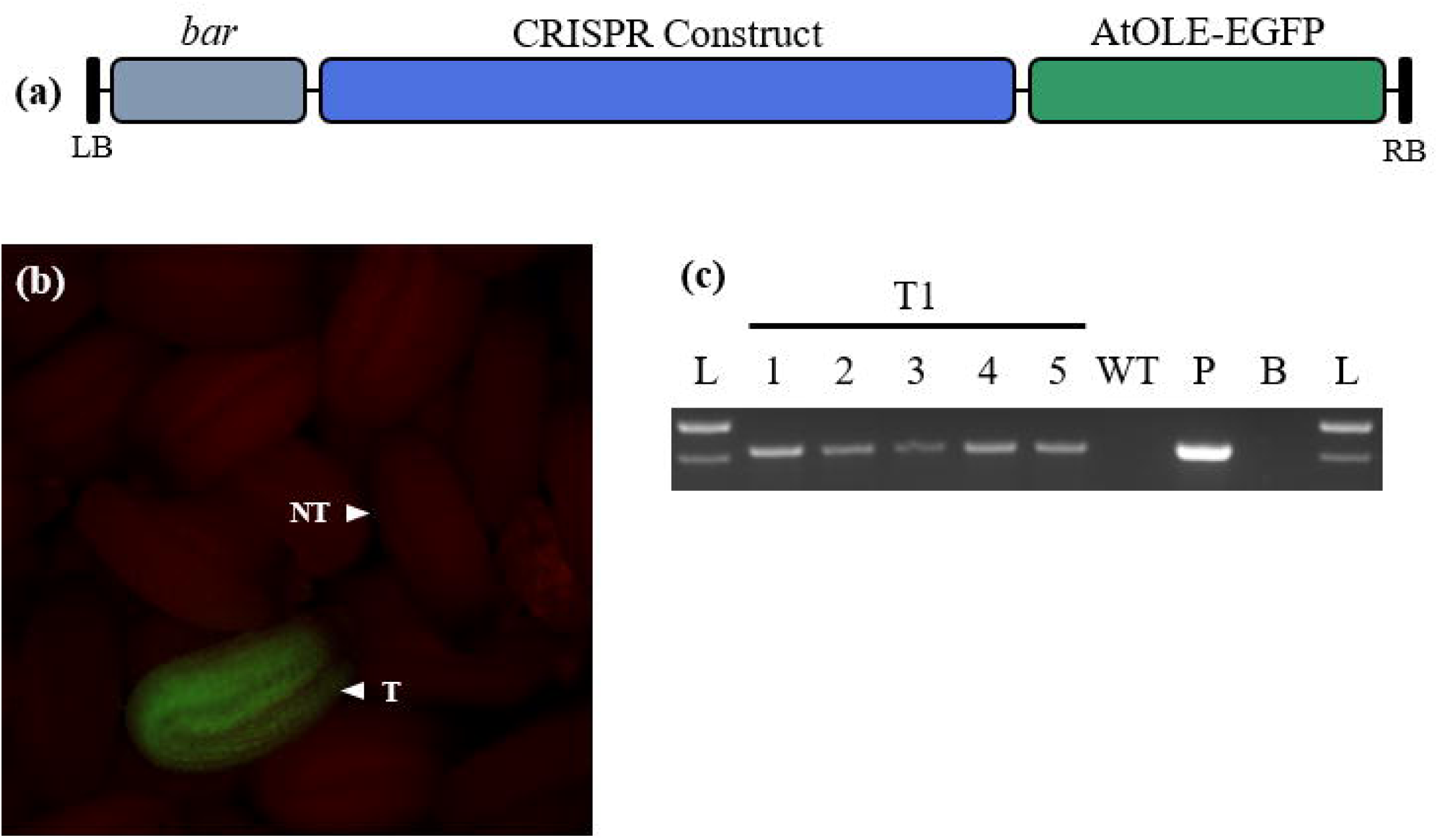
*Capsella grandiflora* floral dip transformation. (a) T-DNA cassette introduced into *C. grandiflora* contains the *AtOleosin* promoter and coding region fused to *EGFP*, a Basta resistance gene (*bar*), and a CRISPR construct. (b) EGFP-expressing mature T1 seeds visualized under a blue fluorescent filter. T: transgenic seed; NT: non-transgenic seed. (c) PCR targeting *bar* in seedlings from EGFP expressing seeds (T1 1-5), a wild type control (WT), the original plasmid (P), a no-template control (B), and ladder (L).

The Arabidopsis floral dip method was adapted for transformation of *C. grandiflora* [24]. Developing fruits and open flowers were removed one day before dipping. Floral meristems were dipped into the infiltration medium with resuspended Agrobacterium for ten seconds, then transferred to a dark, humid container for 24 hours. This procedure was repeated a total of six times. The first four dips were performed every four days, followed by a seven day resting period before two final dips were performed seven days apart.

Importantly, as an obligate outcrosser *C. grandiflora* must be manually pollinated from another individual with a different S-locus haplotype [25]. To meet this requirement the dipped *C. grandiflora* plants were hand-pollinated daily to ensure fertilization of the potential transformed ovules. In Arabidopsis the female reproductive tissue is the primary target of *Agrobacterium*-mediated floral-dip transformation [26], however the effect of Agrobacterium infection on pollen viability in *C. grandiflora* was not known. To ensure viable pollen were present, we hand-pollinated the dipped *C. grandiflora* with untransformed wild type *C. grandiflora* daily using a size 3 round paintbrush. We began hand pollination from the day after the first dip to ten days after the last dip, a total of 38 days. On days the plants were dipped, hand-pollination was performed at least three hours before dipping. The following day, hand-pollination was performed at least three hours after removal from the dark. This allowed to the flowers to dry prior to hand-pollination. The dipping procedure was performed on two sets of either three or four *C. grandiflora* plants.

Seeds were harvested and vacuum desiccated for seven days before storage. Seeds were examined using a stereomicroscope with a blue fluorescent filter to visualize the EGFP (Figure 1B, Zeiss Axiozoom 16). Several EGFP positive seeds were selected for propagation. PCR targeting the Basta resistance gene (*bar)*, was used to verify transgene insertion using wild type leaf tissue as a negative control and the insertion vector as a positive control (Figure 1C). DNA was purified from leaf tissue using phenol/chloroform extraction. All of the propagated EGFP positive seeds were confirmed with PCR to have the T-DNA insert.

Transformation efficiency was calculated by first determining the total number of seeds produced from plants in both transformation attempts. In brief, we determined the average weight of two sets of 1000 seeds from each transformation attempt and then estimated the total number of seeds produced by weighing all of the seeds from plants in each attempt. EGFP positive seeds from each attempt were counted and expressed as the percentage of total seed produced in that attempt giving efficiencies of 0.17 % (39 / 22,667) and 0.52 % (79 / 15,158).

Due to the self-incompatibility of *C. grandiflora* it was necessary to pollinate positive transformants with compatible wild type plants to maintain the transgenes. S-loci haplotypes are diverse within *C. grandiflora* populations [25] making it important to determine which haplotypes are present in any transformed *C. grandiflora* individuals. We used generic S-locus primers SLGF and SLGR from Charlesworth et al. [27] to amplify the S-locus glycoprotein (SLG) from ten individuals, then cloned the PCR product into a TOPO-TA vector (Thermo Fisher 450071). Three colonies were sequenced from each individual to determine the variety of haplotypes in our 83.17 *C. grandiflora* population. We used the sequencing data to design haplotype specific primers for the five distinct alleles present in our population. These primers were used to select mating pairs with different S-loci alleles, thereby allowing us to establish populations of T-DNA transformed lines through breeding. We recommend empirically determining the population variation in SI loci using the approach we have outlined since it may differ from ours depending on the seeds used to establish the population.

With the ability to easily transform *C. rubella* [28], and now *C. grandiflora* via floral dip, we have gained a valuable evolutionary resource. Reverse genetics can now be used to study selfing syndrome within the *Capsella* genus.

## Acknowledgements

We thank the members of the Beilstein lab for help with plant cultivation, tissue collection, and moral support.

